# Microbiome-derived metabolite effects on intestinal barrier integrity and immune cell response to infection

**DOI:** 10.1101/2024.02.06.579072

**Authors:** Lauren Adams, Xiang Li, Richard Burchmore, Richard Goodwin, Daniel M. Wall

## Abstract

The gut microbiota exerts a significant influence on human health and disease. While compositional changes in the gut microbiota in specific diseases can easily be determined, we lack a detailed mechanistic understanding of how these changes exert effects at the cellular level. However, the putative local and systemic effects on human physiology that are attributed to the gut microbiota are clearly being mediated through molecular communication. Here we determined the effects of a number of gut microbiome-derived metabolites on the first line of defence in the gut. Using *in vitro* models of intestinal barrier integrity, and studying the interaction of macrophages with pathogenic and non-pathogenic bacteria, we could ascertain the influence of these metabolites at the cellular level at physiologically relevant concentrations. Many metabolites exerted competing influences on intestinal epithelial or immune cells, specific metabolite effects were noted on barrier function, polarised cytokine release and the lifespan of metabolite treated cells. Our findings reiterate the complexity of understanding microbiome effects on host physiology with many metabolites having contrasting effects on host cells. However, our results underline that microbiome metabolites are crucial mediators of barrier function and the innate response to infection. Understanding the effects of these metabolites at the cellular level will allow us to move towards a better mechanistic understanding of microbiome influence over host physiology, a crucial step in advancing microbiome research.

## Introduction

The gut microbiome, comprising the collection of symbiotic, commensal and pathobiont microorganisms, is of increasing interest due the realisation of the importance of microbes to human health and disease (1). These microbes play a fundamental role in several aspects of human physiology including immune cell development, immune homeostasis, food digestion, enteric nerve regulation and angiogenesis (2, 3). The gut microbiota is shaped by a variety of factors including host genetics, diet, mode of birth and by insults such as antibiotic use and bacterial infection (4, 5). Microbial perturbations are suggested to contribute to many diseases, yet direct causative links remain for the most part elusive and it is unclear in most cases whether microbes play a central role as drivers of disease or whether their appearance is a consequence of disease (6, 7).

Commensal microbes present in the intestinal mucus layer ensure resistance to pathogens by producing antimicrobial substances, modulating the luminal pH, and competing for nutrients (8). Studies in germ-free animals have shown that commensal bacteria are responsible for promoting angiogenesis and development of gut-associated lymphoid tissue and the intestinal epithelium (9). Therefore, commensal microbes play a dual role, exerting protection against pathogens whilst also contributing to formation of a functional intestinal mucosal barrier (9, 10).

However, in diseases such as inflammatory bowel disease (IBD) where patients have an increased gut permeability, particularly during active disease, it is unclear whether an impaired barrier function is the result, or cause, of intestinal inflammation (11, 12). However, it is clear that this disruption in mucosal barrier function can result in the translocation of intestinal microbiota and potentiation of the immune system (13). During intestinal inflammation there are distinct changes in the mucus layer environment which includes decreased mucin production via goblet cells, decreased glycosylation products and a reduction in antimicrobial factors, weakening the protective functionality (14, 15). Moreover, tight junction (TJ) proteins are disrupted during IBD pathogenesis, undermining barrier integrity and contributing to the translocation of microbes (16). Colonising germ-free mice with specific gut microbiota such as *Bacteroides thetaiotaomicron* was shown to counteract this, increasing the expression of genes that encode proteins such as zonula occluden-1 (ZO-1) thus repairing TJs (17). Other microbes, such as commensal *Escherichia coli* C25, have been found to have similar effects, altering the localisation of another TJ protein, claudin-1, and additionally stimulating the secretion of interleukin-8 (IL-8) from intestinal epithelial cells (IECs) (18). Contrastingly hydrogen sulphide producing bacteria, such as *Atopobium parvulum,* which are increased in the intestine of Crohn’s disease (CD) patients with severe inflammation, result in mitochondrial damage in IECs leading to dysfunction and inflammation (19). Therefore, it has been suggested that dysbiosis in the inflamed intestine exacerbates this increased permeability in the gut via a variety of mechanisms and clearly there is complex interplay between the gut microbiota which directly impacts barrier stability in the intestine (17, 18). However, the molecular mechanisms underlying these changes are poorly understood.

Dysfunctional interactions between the intestinal microbiota and the mucosal immune system can also promote a loss of immune tolerance and thus inflammation (20). In IBD large numbers of abundant *Proteobacteria* pass through the mucosal barrier and pathogen-associated molecular patterns (PAMPs), lipopolysaccharide (LPS) and flagellin on the bacterial surface are recognised by toll-like receptors (TLRs) of the innate immune response (21). When the intestinal barrier is disrupted, macrophages and dendritic cells (DCs) sense pathogen associated molecule patterns (PAMPs) with pathogen recognition receptors (PRRs), signalling a downstream activation of central immune response pathways: NF-κB, mitogen activated protein kinases (MAPKs), and interferon regulatory factors (IRFs) (22). The activation of these pathways results in the production of proinflammatory cytokines such as IL-1, IL-6 and tumour necrosis factor alpha (TNF-α), chemokines, and antimicrobial peptides (AMPs) (22, 23). These recruit neutrophils, activate macrophages, and result in dendritic cell maturation, promoting the induction of the adaptive immune response (24, 25). IBD patients have elevated levels of many proinflammatory cytokines in serum and mucosal tissue; thus, it has been postulated that this elevation is primarily due to the uncontrolled immune response to bacterial antigens (20). Therefore, microbiome mediated innate immune system activation contributes to IBD pathogenesis by inducing an inappropriate proinflammatory response.

Microbes in the gut benefit the host by providing defence against pathogens, promote immune maturation and synthesise nutrients while microbe-host interactions involved in disease pathology are underpinned by small molecules and metabolites (26, 27). The gut microbiota interact and communicate with the host via the production of proteins, lipids and small bioactive metabolites that are critical signalling molecules (28). However, microbial dysbiosis and inflammation in diseases such as CD alters the microbe-host metabolome, yet the molecular mechanisms underlying the contribution of this to disease are still poorly understood (29). Microbial metabolites such as bile acids, SCFAs, tryptophan, TMA and benzoate can be used to discriminate between healthy controls and IBD patients. Further investigation is therefore required to elucidate their biological function in relation to immune homeostasis and IBD pathology. To this end here we determined the effects of a variety of microbiome-derived metabolites on intestinal barrier integrity and immune cell response to infection. Using microbial metabolites L-tryptophan, butyrate, trimethylamine, 3,4-TMAB, 4-TMAP, ursodeoxycholic acid, glycocholic acid and benzoate we tested their effects in isolation on the first line of defence of the immune system, the intestinal epithelial barrier and immune cells. This was done in conjunction with commensal and pathogenic *Escherichia coli* strains to determine any additive or inhibitory effects on the response to microbes. Understanding their effects can give useful insights into how changes in the gut microbiome can lead to alterations in intestinal immune homeostasis.

## Materials and methods

### Bacterial strains and cell culture

The pathogenic adherent invasive *Escherichia coli* (AIEC) strain LF82 and intestinal commensal strain *E. coli* K-12 were cultivated in lysogeny broth (LB) or on LB agar. Bacteria were grown in Roswell Park Memorial Institute 1640 (RPMI) medium supplemented with 3% foetal bovine serum (FBS) and 1% L-glutamine at 37°C and 180 revolutions per minute (rpm) before diluting to give a final multiplicity of infection (MOI) of 100. Supernatants were prepared by cultivating bacteria as described but after reaching an OD_600_ of 0.6, cultures were then transferred to a static incubator for 6 hours at 37°C. Cultures were then centrifuged at 400 g for 5 minutes before supernatants were collected and filtered to remove bacteria. RAW 264.7 macrophages and human intestinal epithelial HCT-8 cells were purchased from the American Type Culture Collection (ATCC). RAW 264.7 macrophages were cultured in RPMI that was supplemented with 10% FBS, 1% L-glutamine and 1% penicillin/streptomycin (P/S). HCT-8 cells were cultured in RPMI supplemented with 10% horse serum (HS), 1% P/S, 1% L-glutamine and 5 mM sodium pyruvate. Cells were incubated at 37°C, 5% CO_2_ and passaged every 2-3 days.

### Microbiome metabolites used in this study

Molecules used in this study were dissolved in water to make stocks and stored as aliquots at -20°C. Molecules were fully defrosted before use and diluted to reach desired working concentrations (Table 1).

**Table 1:** List of microbiome-derived metabolites and their concentrations used in this study. The source of the metabolites is also indicated.

| Molecule | Low concentration | High concentration | Catalog | Company |
| --- | --- | --- | --- | --- |
| L-Tryptophan | 10 µM | 30 µM | T0254 | Sigma-Aldrich |
| Sodium butyrate | 0.5 mM | 2 mM | B5887 | Sigma-Aldrich |
| Trimethylamine hydrochloride (TMA) | 10 µM | 30 µM | T72761 | Sigma-Aldrich |
| 3-methyl-4-(trimethylammonio)butanoate (3, 4-TMAB) | 20 µM | 1 mM | custom made | AstraZeneca |
| 4-(trimethylammonio)pentanoate (4-TMAP) | 20 µM | 1 mM | custom made | AstraZeneca |
| Ursodeoxycholic acid (UDCA) | 10 µM | 30 µM | U5127 | Sigma-Aldrich |
| Glycocholic acid hydrate (GCA) | 250 nM | 500 nM | G2878 | Sigma-Aldrich |
| Sodium benzoate | 10 µM | 50 µM | B3420 | Sigma-Aldrich |

### E. coli LF82 growth and survival in RAW 264.7 macrophages and TNF-***α*** production

RAW 264.7 macrophages were seeded at a density of 2 x 10L cells in RPMI supplemented with 3% FBS and 1% L-glutamine, and incubated at 37°C, 5% CO_₂_ for 5-6 hours. Cells were then activated with 100 ng/ml of LPS and supplemented with the appropriate metabolite concentration before incubating for a further 18 hours. Bacterial uptake (MOI of 100) was allowed to proceed for 1 h at 37°C, 5% CO_2_. To determine the extent of bacterial uptake and proliferation, extracellular bacteria were washed away, and 50 μg/ml gentamycin sulphate was added for 1 h to kill any remaining cell-adherent bacteria. RPMI supplemented with 3% FBS, 1% L-glutamine was then added with the appropriate metabolites. Cells were incubated for a further 24 to 72 h. Cells were then harvested before lysing with 2% Triton X-100. Total bacteria were enumerated by counting colony forming units (CFUs) after overnight incubation at 37°C on LB agar plates. TNF-α production was quantified in the supernatant of RAW 264.7 cells using a sandwich ELISA Max Deluxe Set (Biolegend) according to the manufacturers protocol.

### Transepithelial electrical resistance (TEER) and polarized secretion of pro-inflammatory cytokines

The surface of 0.3 cm^2^, 3.0 μm pore sized transwell inserts were coated with 50 μg/ml rat tail collagen. After collagen coating, 1 ml of culture media (RPMI supplemented with 10% horse serum (HS), 1% P/S, 1% L-glutamine and 5 mM sodium pyruvate) was added to the basolateral side of a 24-well tissue culture plate. HCT-8 cells were seeded at a density of 2 x 10D cells in 200 μl of media per insert and incubated at 37°C, 5% CO_2_. TEER was measured using a voltmeter in triplicate for each well. Cells were grown until a monolayer had been achieved represented by TEER value above 300 Ω. TEER was calculated by multiplying the surface area of the transwell (in cm^2^) by the NET resistance (which is the resistance measured minus the resistance of a blank transwell covered by cell culture media). Microbiome metabolites were then added at the chosen concentrations and TEER was measured over a time course of 24 hours. Apical and basolateral supernatants were collected and stored at -80°C for further use. The concentration of IL-6, 1L-8 and IL-15 were quantified using individual ELISA kits from (Biolegend), following the manufacturer’s protocol. Cell lysates were obtained by transferring the transwell inserts into a 24-well plate containing 0.1% TritonX-100 to lyse the cells before lysates were stored at -80°C for future use.

### Quantification of tight junction protein by Western blot

HCT-8 cells were seeded at a density of 2 x 10D cells/ml into a 24-well cell culture plate and metabolites were added as indicated. Cells were incubated at 37°C, 5% CO_2_ for 48 h before infection with LF82 at an MOI of 100 for 3 h. Cells were washed with ice cold phosphate buffered saline (PBS) twice and subsequently lysed for 10 minutes with radioimmunoprecipitation assay lysis buffer (RIPA), supplemented with protease inhibitors. Lysates were frozen at -80°C prior to use. Tight junction and control proteins were detected using antibodies for ZO-1 (catalog number 339100, ThermoFisher Scientific), GAPDH (catalog number 2118, Cell Signalling Technology) and a HRP-conjugated rabbit IgG secondary antibody (catalog number 31460, ThermoFisher Scientific). Membranes were developed by applying an enhanced chemiluminescence agent and imaged with a blot scanner. Blots were set up into biological triplicates and bands were analysed using ImageJ software before one-way ANOVA was performed on Graphpad prism.

### Quantification of caspase-3/7 activity in HCT-8 cells

HCT-8 cells were seeded at a density of 2 x 10□ cells/ml into a 24-well cell culture plate and metabolites were added. Cells were incubated at 37°C, 5% CO_₂_ for 24 h before supernatants were collected. Cells were then lysed using 0.1% TritonX-100 and lysates stored at -80°C. Caspase 3/7 activity was then quantified using the Apo-One Homogenous Caspase-3/7 assay (Promega).

### TMRE-Mitochondrial membrane potential analysis

Mitochondrial membrane potential was measured with a TMRE (tetramethylrhodamine ethyl ester) assay (abcam). HCT-8 cells were seeded at a density of 2 x 10D cells/ml into a 96-well cell culture plate and metabolites were added. Cells were the incubated for 24 h at 37°C, 5% CO_2_. The positive control carbonyl cyanide-p-trifluoromethoxyphenylhydrazone (FCCP), an uncoupler of mitochondrial oxidative phosphorylation, was applied at a final concentration of 20 μM for 10 min before TMRE treatment. The cells were incubated with 1 μM TMRE for 30 min at 37°C, 5% CO_2_, followed by washing twice with 100 μl of PBS containing 0.2% bovine serum albumin (BSA). A volume of 100 μl of PBS containing 0.2% BSA was added to each well, and the fluorescence was measured in a FLUOstar Galaxy plate reader (BMG Labtech) with excitation/emission: 544/590 nm. PBS containing 0.2% BSA was then removed from cells and replaced with 20 μl 0.1% TritonX-100 and cells were kept on ice before storing at -20°C and protein concentration was determined using a BCA assay. Data was normalised using sample protein concentration and shown as TMRE fluorescence percentage of untreated cells. One-way ANOVA was performed using Graphpad prism (versus the control condition - cells without metabolites).

### Quantification of LDH release as a feature of apoptosis

HCT-8 cells were seeded at a density of 2 x 10D cells/ml into a 24-well cell culture plate and the appropriate metabolites were added. Cells were incubated at 37°C, 5% CO_2_ for 24 h before supernatants were collected and LDH levels determined using an LDH-Cytotoxicity Assay Kit (abcam).

## Results

### Small molecule effects on intestinal barrier function

Infection with LF82 resulted in a significant decrease in ZO-1 levels (2.21-fold) in HCT-8 cells compared to uninfected controls (Fig. 1). However, the pre-treatment of cells with specific metabolites prior to LF82 infection reversed this, with L-tryptophan (2.57-fold, *p=*0.0028), butyrate (3.10-fold, *p=*0.0007), TMA (2.42-fold, *p=*0.0067), UDCA (2.19-fold, *p=*0.0001), and benzoate (2.55-fold, *p=*0.0027) each significantly increasing ZO-1 levels compared to infected cells without pre-treatment (Fig. 1B). 3,4-TMAB at 20 μM significantly increased ZO-1 levels (1.8-fold, *p=*0.0071) compared to infection alone, but at a concentration of 1 mM levels of ZO-1 reverted to those of infection without pre-treatment suggesting 3,4-TMAB may be protective, but only at lower physiologically relevant concentrations (30). The pre-treatment of cells with 4-TMAP which is highly similar in structure to 3,4-TMAB, significantly increased ZO-1 (2.76-fold, *p*<0.0001) expression compared to infection alone when administered at 20 μM. However, unlike 3,4-TMAB when the concentration was further increased to 1 mM, ZO-1 levels were even further increased (1.3-fold, *p=*0.0405).

**Figure 1.**
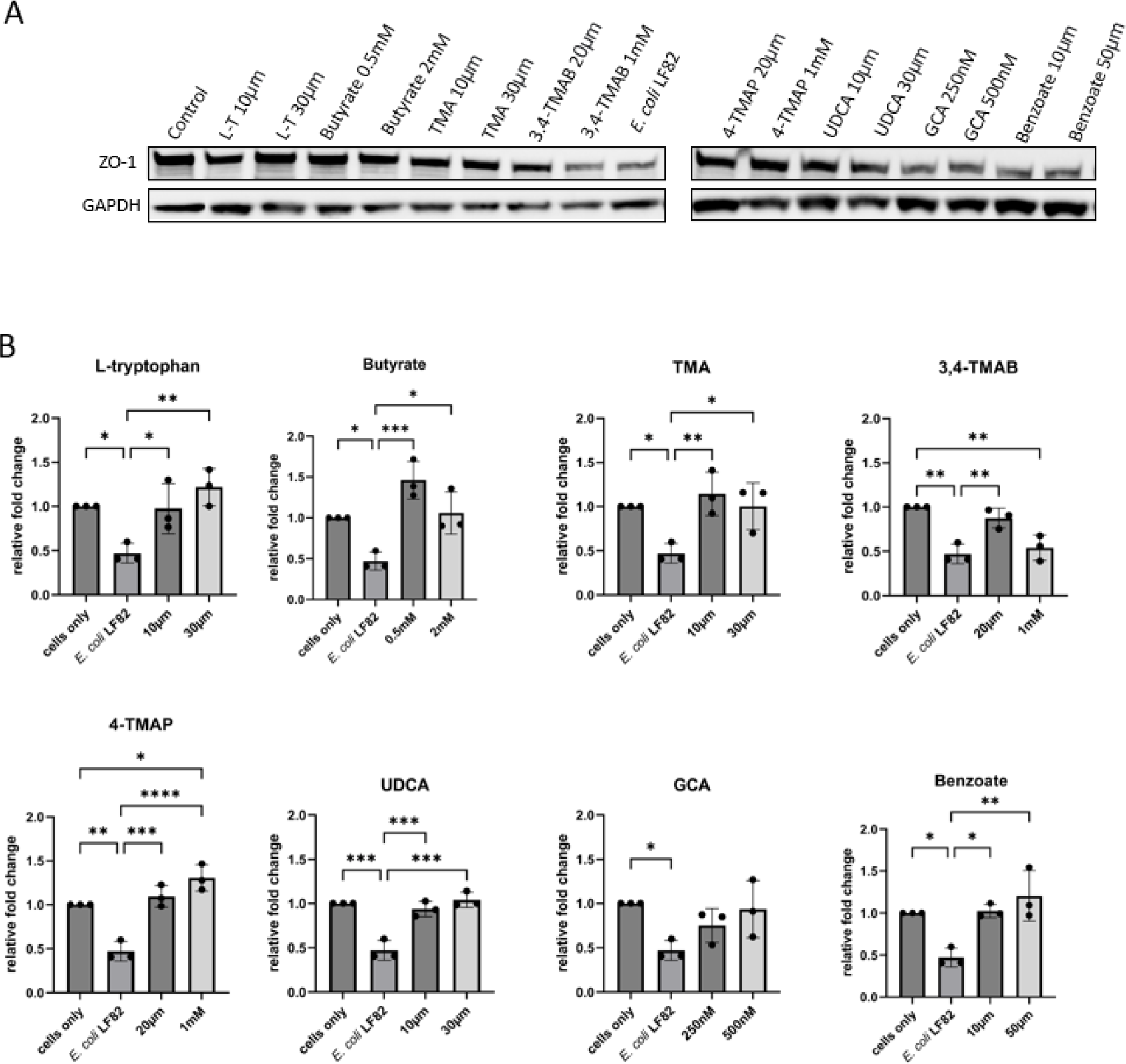
HCT-8 exposure to metabolites rescues ZO-1 expression following infection with *E. coli* LF82. A) Image shows ZO-1 and GAPDH expression (Lanes 1-18) in HCT-8 cells. Cells were pre-treated for 48 h with metabolites at different concentrations as indicated, followed by infection with *E. coli* LF82 for 3h. Lane 1 shows HCT-8 cells without pre-treatment or infection and Lane 10 shows infected HCT-8 without pre-treatment with any metabolite. B) Graphs are labelled with the molecule that was used to pre-treat HCT-8 cells before infection and show the relative fold change in ZO-1 levels compared to control (HCT-8 cells without treatment or infection). Data shown represents three biological replicates and ImageJ was used for analysis. One-way ANOVA was performed to test significance by comparing all conditions using Graphpad prism. *p<0.05, **p<0.01, ***p<0.001, ****p<0.0001 versus the control condition (infection without metabolites) was considered statistically significant.

To investigate the repercussions of these changes in ZO-1 expression for barrier function, an *in vitro* model to test transepithelial electrical resistance (TEER) was employed. A HCT-8 cell monolayer was grown on transwell inserts (TEER value above 300 Ω) before metabolites were added. *E. coli* supernatants were used as controls and upon addition of bacterial supernatants or the metabolites for testing into the apical compartment, the temporal changes in barrier function were measured. The control *E. coli* K-12 supernatant significantly increased barrier function 0.77-fold (*p=*0.0376) and 0.74-fold (*p=*0.0223) after 3 h and 6 h, respectively, compared to the control (Fig. 2). However, only the addition of 2 mM butyrate significantly altered barrier function, increasing relative TEER 0.7-fold (*p=*0.0058) after 6 h, compared to HCT-8 only control. While a number of other metabolites altered TEER none of these changes were significant.

**Figure 2.**
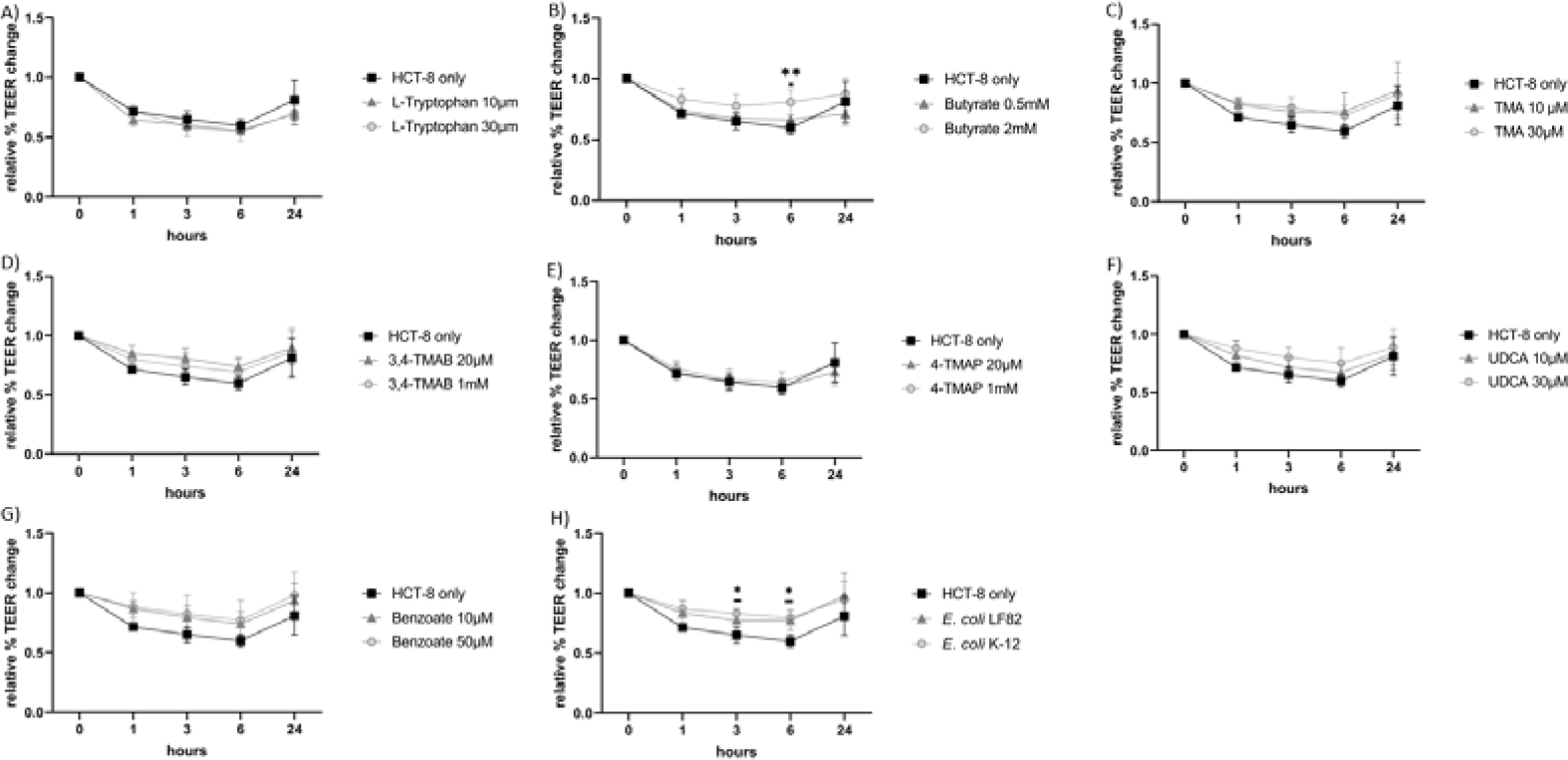
TEER as a model for HCT-8 barrier function in response to metabolites. HCT-8 cell monolayers were treated with indicated metabolites or control supernatants from *E. coli* LF82 and *E. coli* K-12. TEER was measured over a 24 h period and results are shown as a relative percentage change compared to control (0 h TEER reading). Data shown is the mean of three biological replicates ± standard deviation (SD) (error bars). Two-way ANOVA was performed to test significance between untreated and monolayers treated with different molecule concentrations at each time point. \**p*<0.05, \*\**p*<0.01, \*\*\**p*<0.001, \*\*\*\**p*<0.0001 was considered statistically significant.

### Microbiome metabolites as cytotoxic stressors, inducers of apoptosis and inhibitors of mitochondrial function

As barrier integrity is also influenced by lifespan and health of IECs we next investigated if cytotoxicity of metabolites or their induction of cell death was playing a role. LDH is a stable cytoplasmic enzyme found in all cells and its release here was used to determine whether the microbiome metabolites were potentially cytotoxic to HCT-8 cells. Supernatants from both *E. coli* strains were not cytotoxic to HCT-8 cells but 2 mM butyrate significantly increased cytotoxicity (1.98-fold, *p=*0.0320), whereas 1 mM of 4-TMAP significantly decreased cytotoxicity (4.7-fold, *p=*0.0456)(Fig. 3). Similarly, treating HCT-8 cells with 2 mM butyrate significantly increased caspase-3/7 activity (7.89-fold, *p*<0.001) indicating progression towards apoptosis (Fig. 4). Given reported communication between microbes and mammalian mitochondria, and the importance of mitochondria in maintaining cell function, we also investigated the effect of the metabolites on mitochondrial activity. A drop in mitochondrial membrane potential is indicative of a drop in mitochondrial activity, which is a negative indicator of overall cell health. HCT-8 cells were exposed to metabolites, or *E. coli* LF82 and *E. coli* K-12 supernatants as controls, for 24 h and a TMRE-mitochondrial membrane potential assay undertaken. TMA (1.62-fold, *p=*0.0018), 4-TMAP (1.6-fold, *p=*0.0005), GCA (1.76-fold, *p=*0.0011) and benzoate (1.70-fold, *p=*0.0127) all significantly impaired mitochondrial activity (Fig. 5). Again, these data point towards these metabolites inducing significant cellular stress. However despite evidence of cytotoxicity and cell stress associated with specific metabolites, this did not translate into undermining of IEC barrier function. Therefore we examined further phenotypes associated with the IEC barrier.

**Figure 3.**
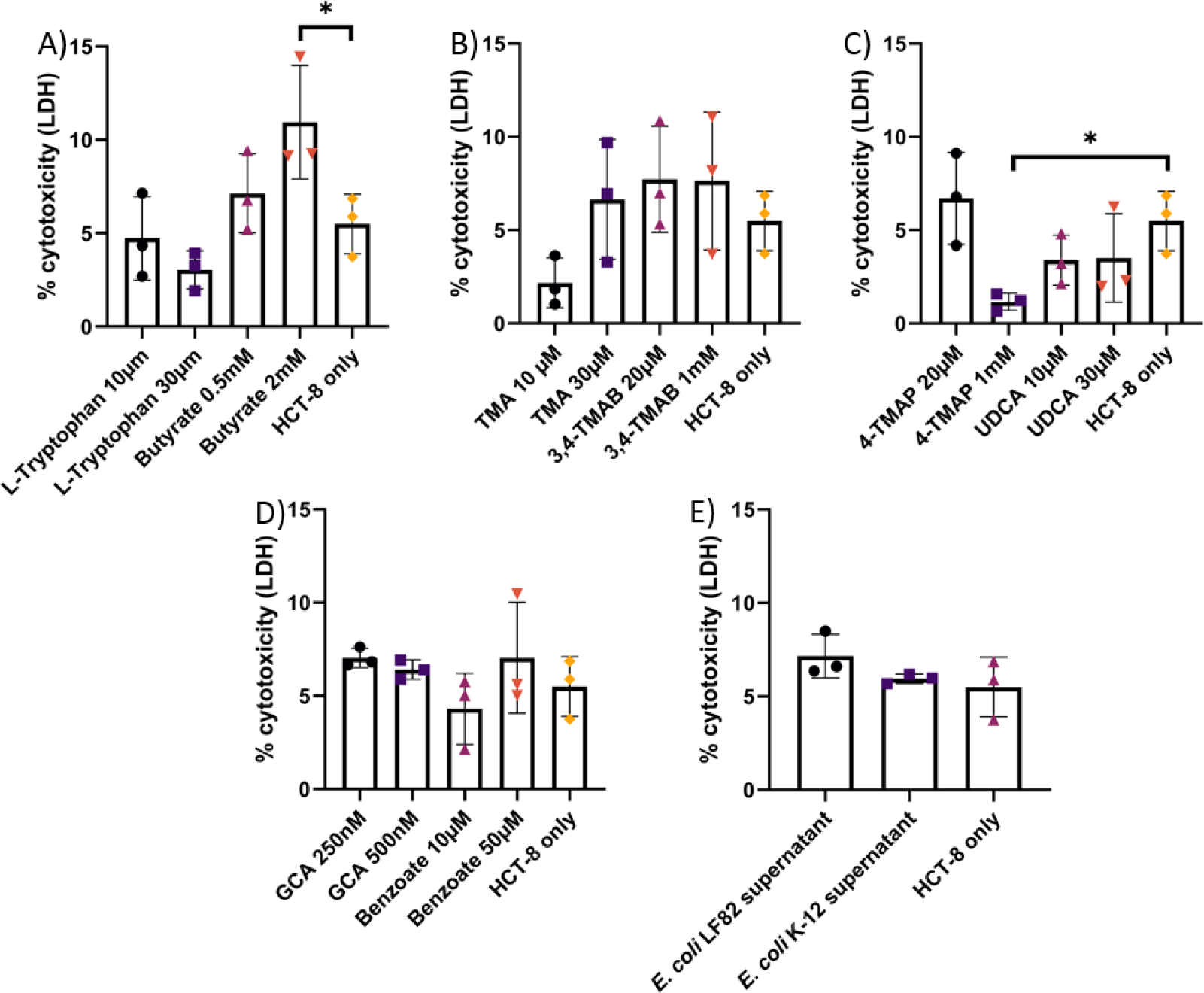
LDH release from HCT-8 cells as an indicator of metabolite cytotoxicity. HCT-8 cells were exposed to metabolites and the supernatants from *E. coli* LF82 and K-12 for 24 h. HCT-8 cells only and cells treated with 2% triton-x were used as low and high LDH release controls, respectively. The percentage of cytotoxicity was calculated as %=[(measured absorbance of sample-low control)/(high control-low control)] x100. Data are shown as the mean of three biological replicates ± standard deviation (SD) (error bars). One-way ANOVA was performed across all metabolites and \**p*<0.05, \*\**p*<0.01, \*\*\**p*<0.001, \*\*\*\**p*<0.0001 versus the control condition (cells without molecules) was considered statistically significant.

**Figure 4.**
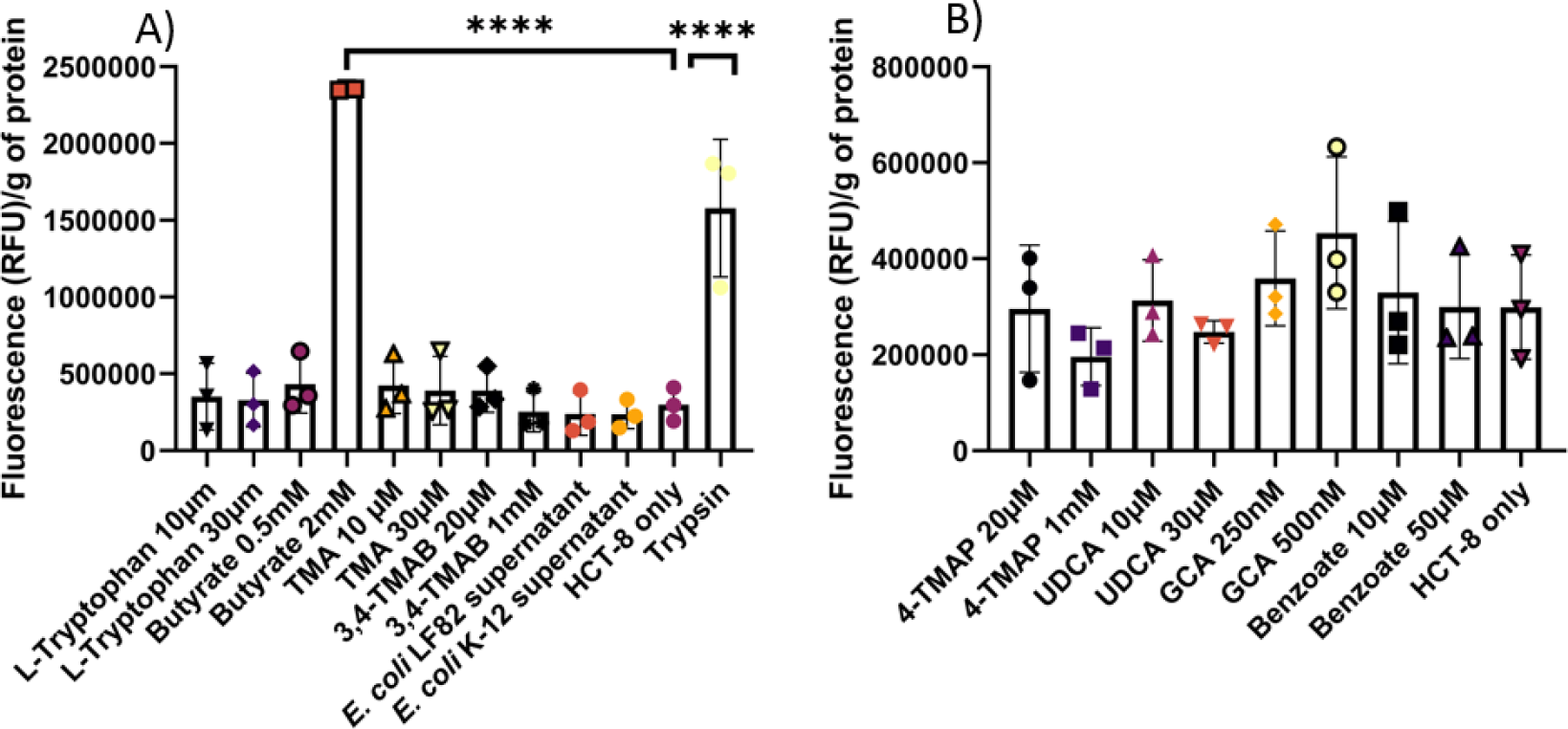
Caspase-3/7 activation in metabolite-treated HCT-8 cells. HCT-8 cells were treated with metabolites and supernatants from *E. coli* LF82 and K-12. Relative fluorescence units (RFU) were normalised to per gram of protein in the cell lysate. Data are shown as the mean of three biological replicates ± standard deviation (SD) (error bars). One-way ANOVA was performed across all metabolites and \**p*<0.05, \*\**p*<0.01, \*\*\**p*<0.001, \*\*\*\**p*<0.0001 versus the control condition (cells without molecules) was considered statistically significant.

**Figure 5.**
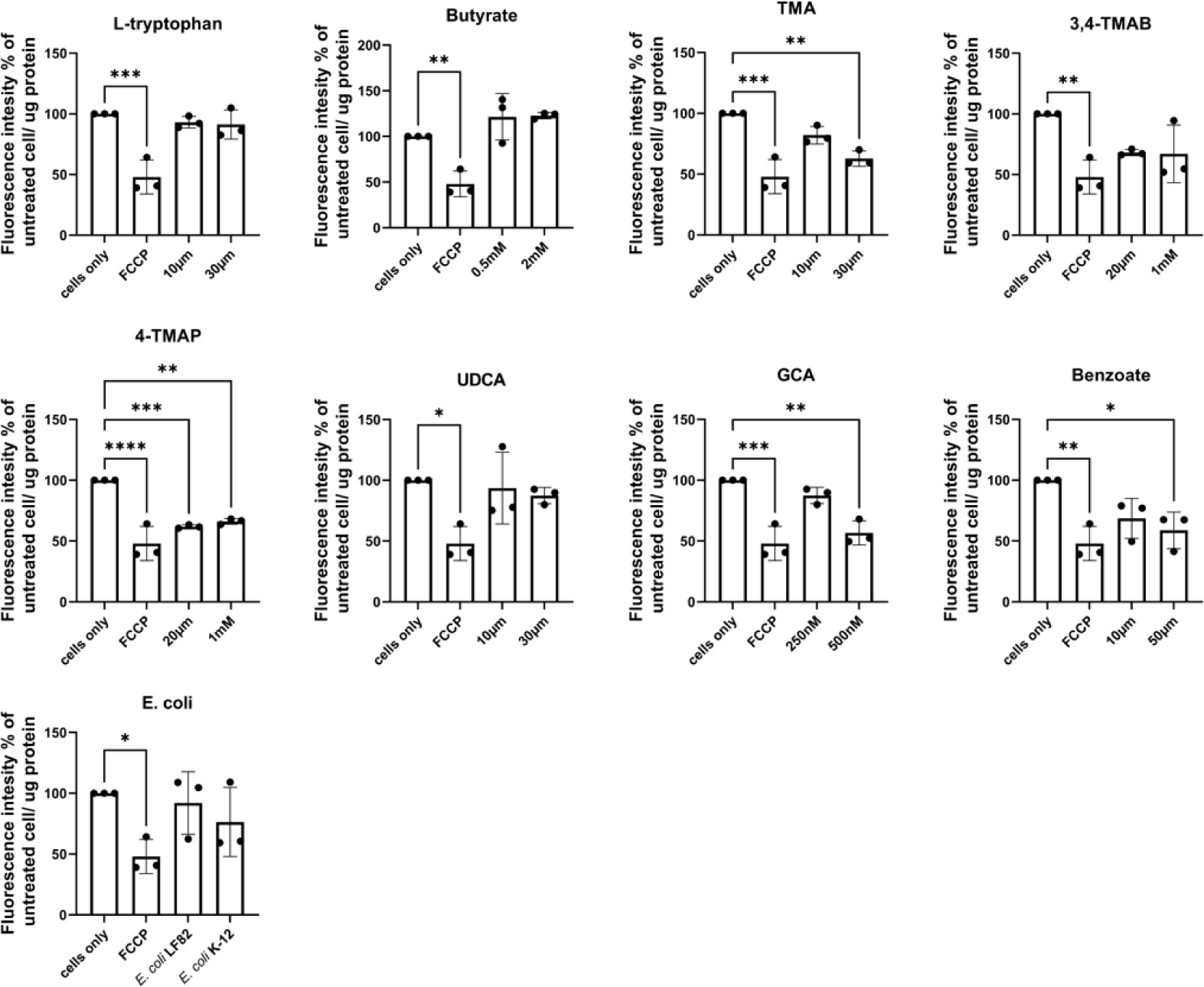
Effect of metabolites on mitochondrial membrane potential. HCT-8 cells were treated with metabolites or bacterial supernatants for 24 h. Cells alone and FCCP were used as high and low mitochondrial membrane potential controls, respectively. Fluorescence was measured and calculated as a percentage of high control (cells only) then normalised to μg of protein in cell lysate. Data are shown as the mean of three biological replicates ± standard deviation (SD) (error bars). One-way ANOVA was performed to compare conditions to control (cells only) and \**p*<0.05, \*\**p*<0.01, \*\*\**p*<0.001, \*\*\*\**p*<0.0001 were considered statistically significant.

### Cytokine secretion from polarized IEC monolayers in response to metabolites

Supernatants were simultaneously collected from the basolateral and apical compartments of the monolayer assay and screened to determine if polarized release of cytokines was occurring in response to apical treatment of the monolayers. UDCA and 3,4-TMAB elicited significantly increased IL-6 release into both the apical and basolateral compartments, compared to the control (Fig. 6). In contrast L-tryptophan, butyrate, TMA, 4-TMAP and benzoate stimulated significant increases in apical but not basolateral secretion of IL-6. The supernatants from *E. coli* LF82 and *E. coli* K-12 elicited no IL-6 release.

**Figure 6.**
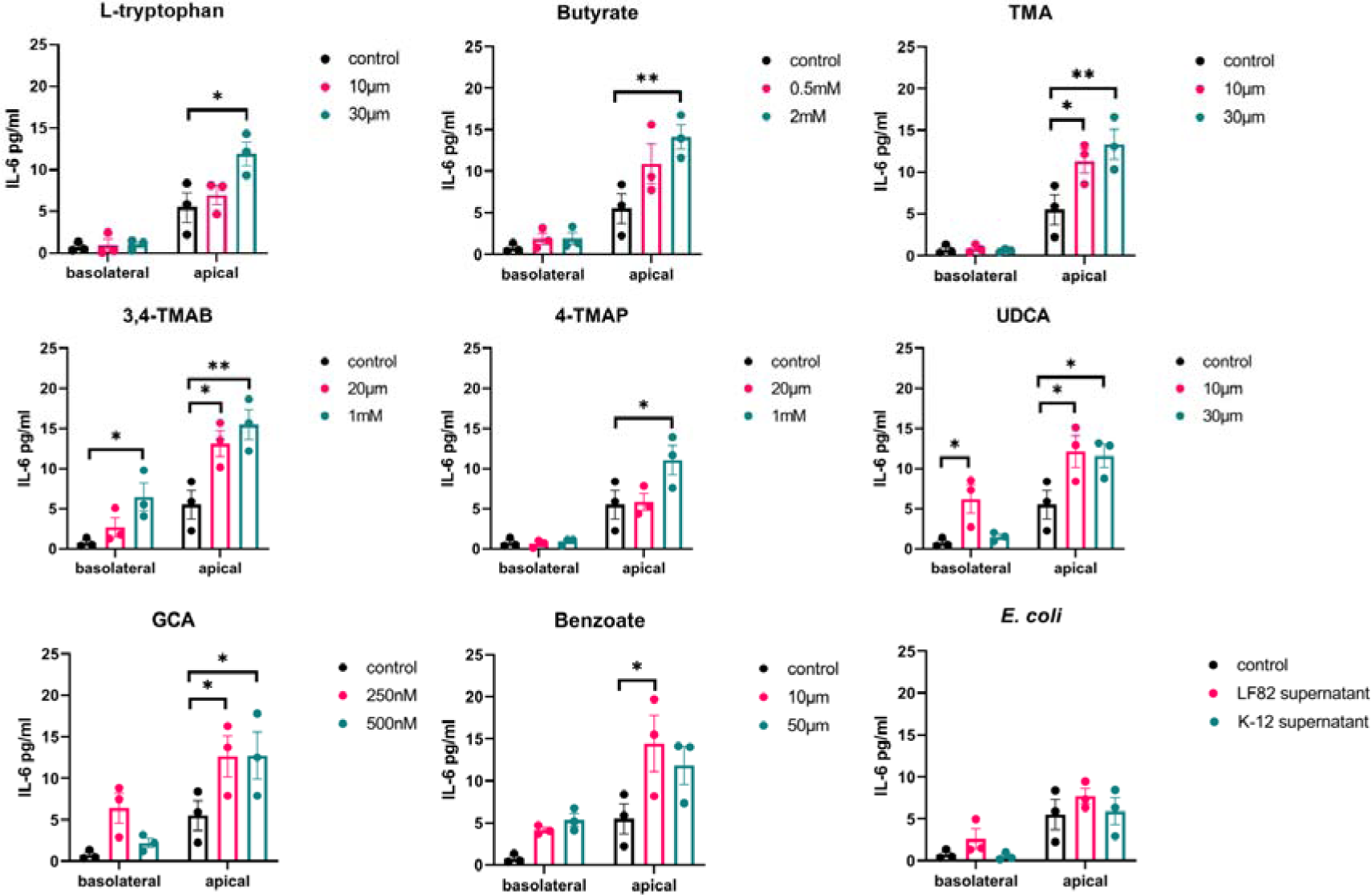
IL-6 release into apical and basolateral epithelial compartments. The apical side of the HCT-8 TEER barrier model was stimulated with supernatant from *E. coli* strains LF82 and K-12 cultures and specific microbiome-derived metabolites. After 24 h the supernatants were collected from the apical and basolateral epithelial compartments, separately. IL-6 was quantified using ELISA and the data is shown as the mean of 3 biological replicates ± standard error of the mean (SEM) (error bars). Two-way ANOVA was performed for each molecule versus the control condition (cells without treatment). *p<0.05, **p<0.01, ***p<0.001, ****p<0.0001 is considered statistically significant.

While the supernatant from the pathogen *E. coli* LF82 significantly increased IL-15 secretion into the basolateral compartment from HCT-8 monolayers (2.2 fold, *p=*0.0075), that of the non-pathogenic *E. coli* K-12 in contrast significantly reduced its secretion (9-fold, *p=*0.0445; Fig. 7). Butyrate (3.2 fold, *p=*0.02) and benzoate (4.12 fold, *p=*0.0003) increased IL-15 basolateral secretion while its apical secretion was significantly increased with L-tryptophan (2.15-fold, *p=*0.034), UDCA (3.95-fold, *p=*0.0014) and with GCA (4.3-fold, *p*<0.0001). 3,4-TMAB and 4-TMAP had the most dramatic effects with 3,4-TMAB significantly increased both apical (2.27 fold, *p=*0.0085) and basolateral (2.8-fold, *p=*0.0437) secretion, while 4-TMAP appeared to strongly induce drive polarized secretion, decreasing basolateral secretion 9-fold (*p=*0.0038) and increasing apical secretion over 4-fold (*p*<0.0001). As expected pathogenic *E. coli* LF82 supernatant elicited a strong IL-8 response both apically (13.1-fold, *p*<0.0001) and basolaterally (8.5-fold, *p=*0.0280) that was completely absent in response to *E. coli* K-12 (Fig. 8). No metabolite tested significantly affected the secretion of IL-8 into either the apical or basolateral compartments. Again these results underlined that when tested in isolation, these metabolites have highly specific and often competing effects on the intestinal immune response.

**Figure 7.**
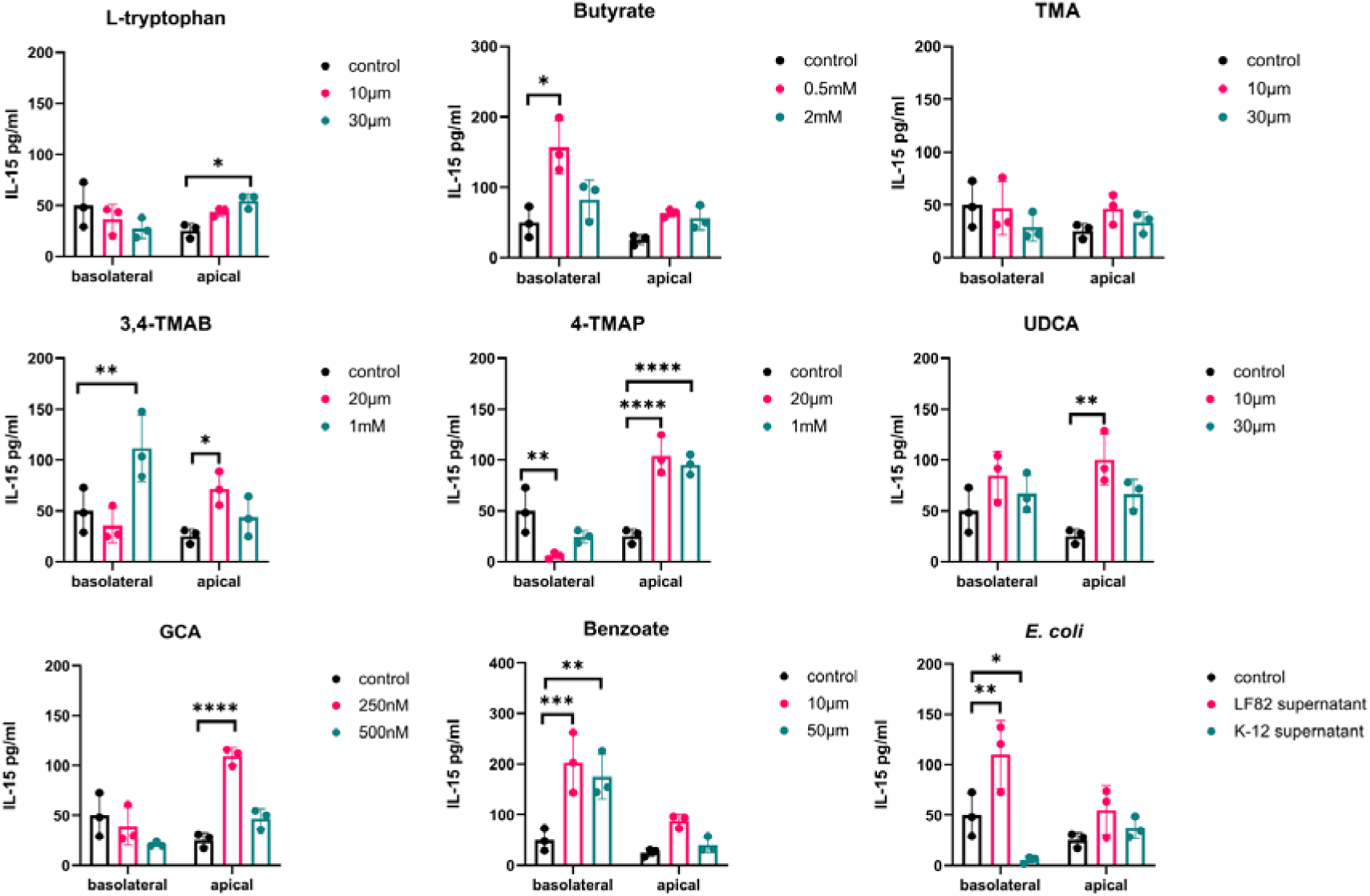
IL-15 release into apical and basolateral epithelial compartments. Apical treatment of HCT-8 monolayers was carried out with named metabolites or bacterial supernatants as controls. After 24 h secreted IL-15 was quantified and the data is shown as the mean of 3 biological replicates ± standard deviation (SD) (error bars). Two-way ANOVA was performed for each molecule versus the control condition (cells without treatment). \**p*<0.05, \*\**p*<0.01, \*\*\**p*<0.001, \*\*\*\**p*<0.0001 was considered statistically significant.

**Figure 8.**
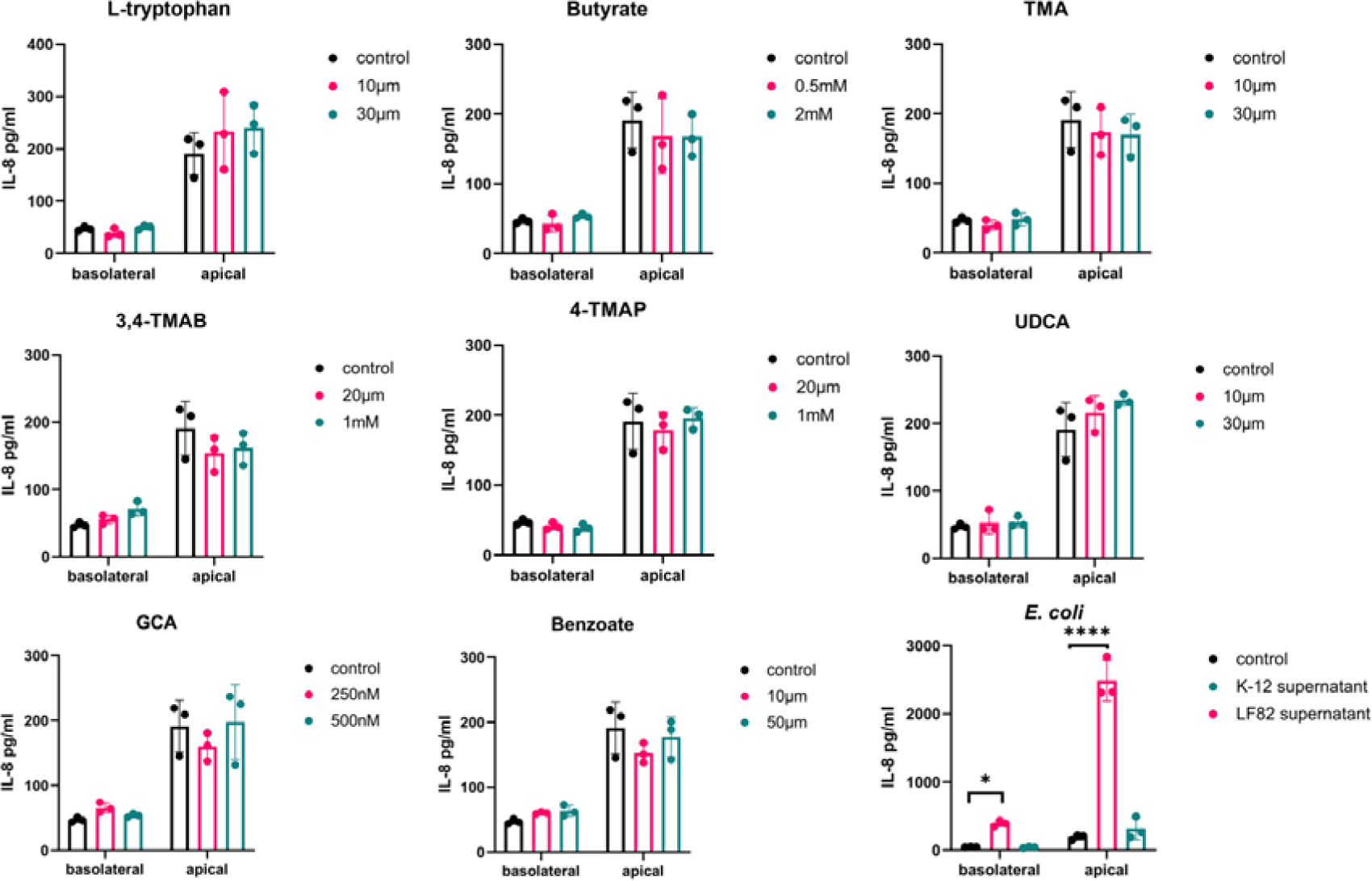
IL-8 release into apical and basolateral epithelial compartments. Apical treatment of HCT-8 monolayers was carried out with named metabolites or bacterial supernatants as controls. After 24 h supernatants were collected from the apical and basolateral epithelial compartments and IL-8 quantified. The data is shown as the mean of 3 biological replicates ± standard deviation (SD) (error bars). Two-way ANOVA was performed for each molecule versus the control condition (cells without treatment). \**p*<0.05, \*\**p*<0.01, \*\*\**p*<0.001, \*\*\*\**p*<0.0001 was considered statistically significant.

### Microbiome metabolites affect macrophage response to E. coli LF82 infection

RAW 264.7 macrophages were stimulated with LPS before treatment with the metabolites at a range of concentrations. Macrophages were then exposed to pathogenic *E. coli* LF82 and uptake allowed to proceed. Cells were lysed 24 h or 72 h post-infection and CFU/ml were calculated. After 24 h no metabolite treatment had a significant effect on bacterial clearance from macrophages (Fig. 9a-b). However, at 72 hpi macrophages treated with the lower concentration of GCA (250 nM) had a significantly increased intracellular bacterial burden (3.3-fold, *p*=0.0167) compared to infection alone (Fig. 9c-d). To rule out this being an artefact due to GCA effects on bacterial growth, a dose response growth curve was undertaken. There were no significant changes in LF82 growth in the presence of GCA (Supplementary Figure 1), indicating GCA increases intracellular bacterial number, likely due to inhibition of clearance of LF82 by RAW 264.7 macrophages.

**Figure 9.**
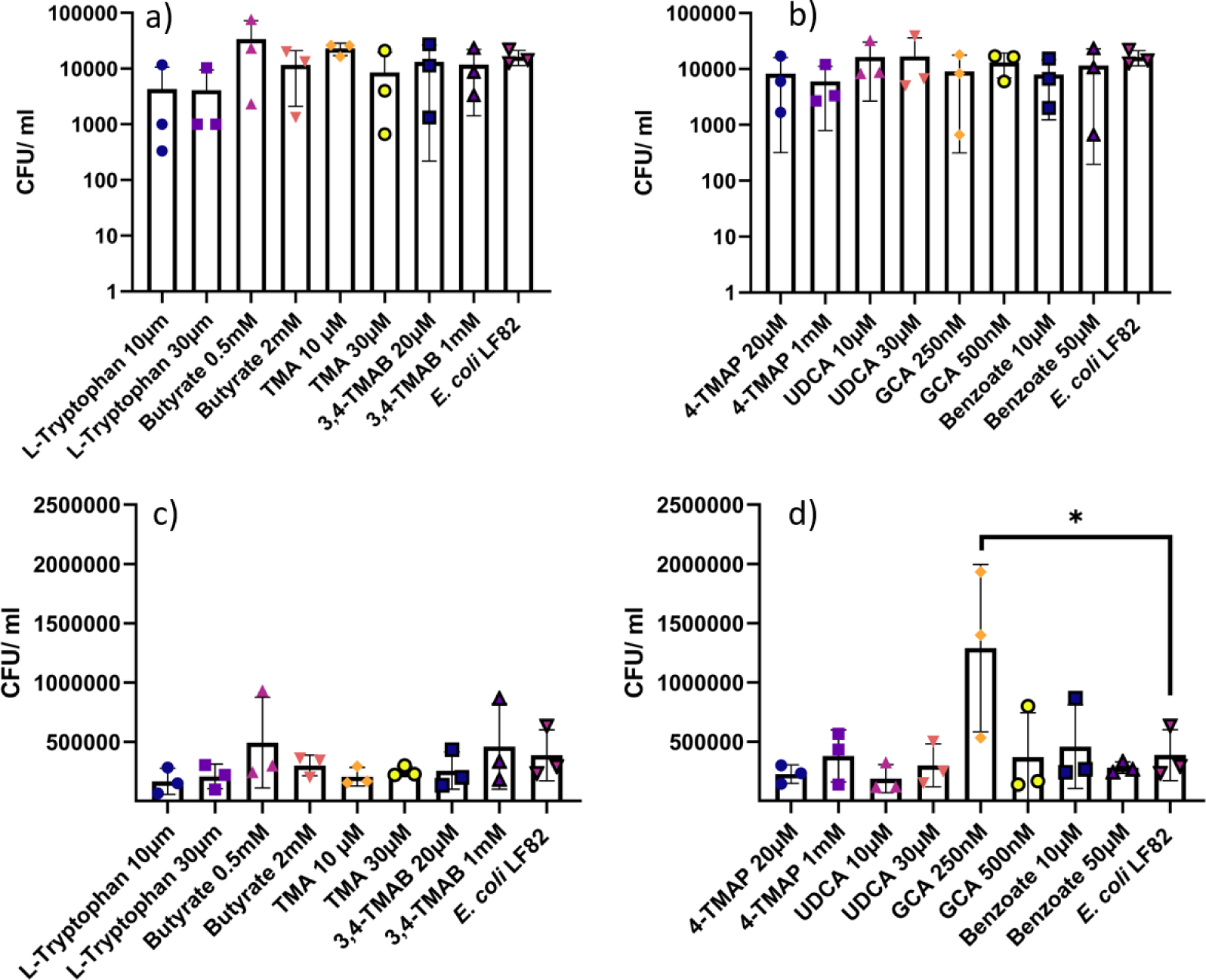
Intracellular *E. coli* LF82 number in RAW 264.7 macrophages after metabolite treatment. RAW 264.7 macropahges were treated wtih various microbial metabolites at different physiologically relevant concentrations before exposure to *E. coli* LF82 and incubation for 24 h (a-b) or 72 h (c-d). Data are shown as the mean of three biological replicates ± standard deviation (SD) (error bars). One-way ANOVA was performed across all metabolites and \**p*<0.05 versus the control condition (infection without metabolites) was considered statistically significant.

The supernatants from RAW 264.7 macrophages from the intracellular bacterial survival assay were collected and used to quantify the secretion of the pro-inflammatory cytokine TNF-α. Given the data in Figure 7 it was hypothesised that only cells pre-treated with GCA would have significantly altered TNF-α levels. However, 24 hours post-infection treatment with 0.5 mM butyrate had significantly increased TNF-α secretion (1.5-fold, *p*=0.0032) (Fig. 10a). At 72 hpi TNF-α release was significantly increased across a range of metabolite treated cells including those treated with; 2 mM butyrate (2.0-fold, *p=*0.0477), 20 μM 3M-4-TMAB (2.1-fold, *p=*0.0251), 250 nM and 500 nM GCA (2.3-fold, *p=*0.0049 & 0.0060 respectively) and 10 μM and 50 μM benzoate (2.4 & 2.1-fold, *p=*0.0021 & 0.0204 respectively)(Fig. 10c-d). This increase in TNF-α secretion indicated that the inflammatory response of RAW 264.7 macrophages was being amplified by the presence of certain metabolites, independent of bacterial burden.

**Figure 10.**
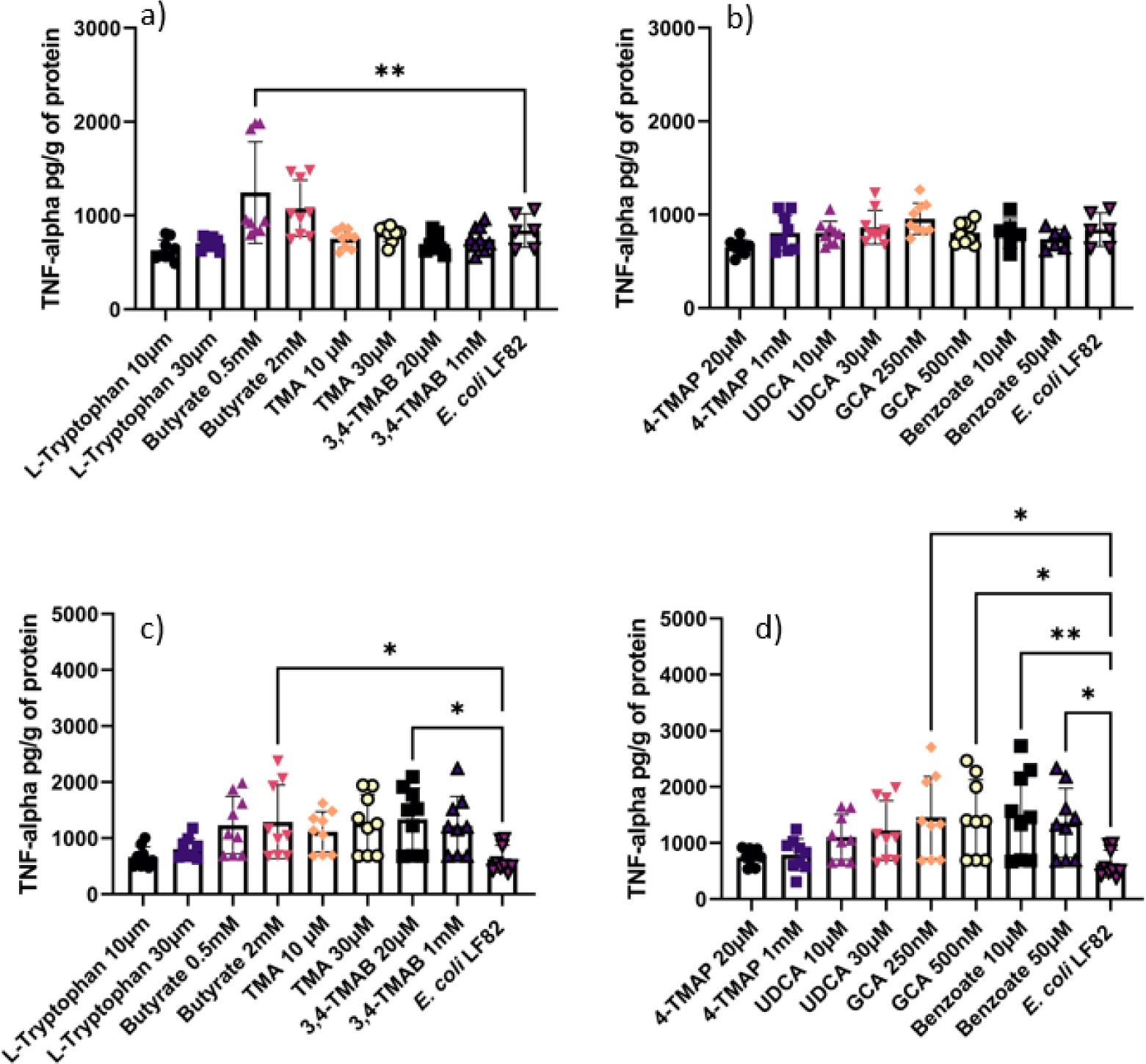
TNF-α released into supernatant of RAW 264.7 macrophages following treatment with metabolites and exposure to *E. coli* LF82. Macrophages were exposed to various microbial metabolites at different physiologically relevant concentrations before allowing uptake of *E. coli* LF82 and infection to proceed for a-b) 24 h and c-d) 72 h. TNF-α was quantified in the cell supernatant and normalised to per gram of protein in cell lysate. Data is shown as the mean of 9 technical replicates ± standard deviation (SD) (error bars). One-way ANOVA was performed across all metabolites and \**p*<0.05, \*\**p*<0.01, \*\*\**p*<0.001, \*\*\*\**p*<0.0001 versus the control condition (infection without metabolites) was considered statistically significant.

## Discussion

It is evident that microbe-host molecular interactions play a role in exerting gut microbiome effects on mammalian and human physiology. However, understanding the mechanism of action of such molecules remains of the upmost importance if the field of microbiome science is to progress. Here taking previously identified microbiome-derived molecules, or by products of bacterial metabolism (metabolites) in the gut, we endeavoured to understand their mechanisms of action at the cellular level. Each was tested at what was deemed a physiologically relevant concentration based on the published literature and they were tested in conjunction with pathogenic and commensal microorganisms.

Studies have shown that in intestinal inflammation as seen in IBD a loss of the tight junction protein ZO-1 occurs that precedes the disruption in barrier integrity leading to further inflammation (31, 32). Furthermore LF82, the pathogen used as a control here, impairs barrier function by decreasing ZO-1 levels (33). Thus, in the first instance it was important to evaluate the impact the microbiome molecules of interest have on ZO-1 expression during infection. L-tryptophan, butyrate, TMA, 4-TMAP, UDCA and benzoate significantly increased ZO-1 levels in the presence of LF82 supernatants compared to cells untreated with supernatants or metabolites. Therefore these molecules were contributing to a protective effect, rescuing ZO-1 levels to those of cells which were not treated with pathogen supernatant. L-tryptophan, butyrate and UDCA have all previously been reported to protect intestinal barrier function, so the effect noted here is not unexpected and could be a broader protective mechanism whereby gut microbiome metabolites stimulate ZO-1 expression to aid in overcoming infection (34–36). TMA and benzoate induction of increased levels of ZO-1 was unexpected as both have been linked to inflammation in the gut (37–39). Yet both molecules have antimicrobial properties and are toxic towards *E. coli* so it is possible that any positive effects on ZO-1 levels seen here were indirect due to the metabolites counteracting the negative effects of the bacterial supernatant treatment (40, 41). Interestingly, both 3,4-TMAB and 4-TMAP had similar effects at low concentrations, increasing ZO-1 levels. Given their similar structure and targeting of fatty acid oxidation it was thought both would exert similar effects in all assays. However, increasing the concentration to 1 mM impaired the protective effect of 3,4-TMAB, and ZO-1 levels were reduced. In contrast 4-TMAP increased ZO-1 levels at a higher concentration. While changes in metabolism can affect ZO-1 expression, the contrasting outcomes here indicate the effects of these metabolites are likely more subtle that simple induction of metabolic shifts and underscore the challenges in teasing apart molecular effects of these metabolites, even in simplistic single cell models as used here (42, 43).

After confirming that the metabolites tested can have a protective or deleterious effect on ZO-1 levels, TEER was used as a direct measurement of barrier function. Treatment of cells with 2 mM butyrate significantly increased barrier function compared to the control. This finding is supported by other studies that have found butyrate to increase the production, and regulate the assembly, of tight junction proteins (44, 45). Given the potentially for cell death to contribute to disruption of the epithelial barrier it was important to confirm whether these molecules of interest were cytotoxic or had the potential to induce cell death through apoptosis or disruption of mitochondrial function. IEC cell death has been directly linked to the development of IBD as IBD patients have been found to have higher levels of cell apoptosis compared to healthy controls (46). The higher concentration of butyrate used was the only molecule to show both a cytotoxic effect and the potential to induce apoptosis. This finding was not unexpected as studies have reported butyrate induction of apoptosis in other epithelial cell lines likely contributing to butyrate driven turnover of intestinal epithelial cells (47, 48). Contrastingly, 4-TMAP significantly reduced LDH release compared to the control, suggesting it exerted a protective effect on IECs. Studies have reported that IECs have decreased levels of ATP when high fat diets shift metabolism towards fatty acid β-oxidation, resulting in cellular death via necrosis (49, 50). As 4-TMAP can inhibit fatty acid oxidation (204), and prevent ATP exhaustion, this may delay necrosis and other forms of cell death (30, 50).

Mitochondrial metabolism and function are known to play an important role in regulating immune cells and maintaining the intestinal epithelial barrier (51). Furthermore, studies have linked dysregulated mitochondrial function in IECs to CD onset and disease severity (52). This study measured mitochondrial membrane potential as an indicator of mitochondrial activity as it is an essential parameter involved in ATP synthesis, respiratory rate, and production of ROS (53). Our study shows that TMA, 4-TMAP, GCA and benzoate can reduce mitochondrial membrane potential at specific concentrations. This finding supports the emerging evidence that microbiome derived small molecules including TMA, 4-TMAP and bile acids alter mitochondrial function (30, 54). The role of TMA in disease has been overlooked due to a focus on the metabolised product, TMAO (37). Therefore, the exact mechanism involved in TMA reduction of mitochondrial membrane potential remains to be fully elucidated and could be an interesting marker of microbial associated mitochondrial dysfunction in disease. Furthermore, 4-TMAP has been found to inhibit enzymes that are involved in carnitine synthesis and fatty acid transportation into the mitochondria (30, 55). This inhibition can lead to a loss of mitochondria function which is observed in this study and may have negative consequences for health (56). Furthermore, studies found specific secondary bile acids contribute to mitochondrial swelling and increase the permeability of the membrane after binding to specific membrane proteins and farnesoid X receptors (57). This triggers mitochondrial fission and results in disordered energy metabolism and apoptosis (57, 58). Therefore, it can be suggested that GCA is another bile acid that impairs mitochondrial function. To date there has been little evidence to suggest benzoate plays a role in mitochondrial dysfunction as seen here, and studies have suggested it may even have a protective effect in neural cells by decreasing mitochondrial caspase-3/7 and ROS (59). As this study shows benzoate impairs mitochondrial membrane potential, it may be that the effect is cell type dependent meaning the high benzoate levels in the intestine may result in cellular damage and inflammation (39, 53, 59).

IECs have two functionally and biochemically different surfaces that play specific roles in cellular function (60). The apical surface faces the intestinal lumen and is mostly involved in absorption and secretion, whereas the basolateral surface mediates the interaction and attachment to underlying neighbouring cells via integrin proteins (61, 62). Furthermore, specific cytokines such as IL-6, IL-15 and IL-8 are secreted at higher levels from the epithelium of the ileum and colon of CD patients (63). These cytokines are pleotropic and have the potential to be proinflammatory under specific conditions (64). Studies have shown that the apical or basolateral secretion of cell signalling molecules including cytokines have different outcomes for immune cell recruitment and activation (65). In particular, the apical release of IL-6 and IL-8 has been associated with increasing neutrophil activation, overproduction of free radicals and skewing macrophage polarization towards a pro-inflammatory M1 phenotype (62, 65). This study found that metabolite treatment of cells did not change IL-8 release from apical or basolateral surfaces. However, LF82 supernatant used as a control significantly increased IL-8 release from both surfaces, but more so apically. Certain pathogenic *E. coli* strains have been shown to induce IL-8 secretion, resulting in reassembly of TJ proteins and increased permeability which allows the transmigration of polymorphonuclear leukocytes (PMNs) to cross epithelium into the lumen (66, 67). This effect has been shown after bacteria accumulate and attach to host cells via the pilus (67, 68). As this study used supernatant instead of bacterial infection, it can be suggested that LF82 can chemically signal IL-8 secretion without using virulence mechanisms which can potentially lead to recruiting PMNs to the intestinal lumen and skew activation towards a pro-inflammatory phenotype. However, it was clear that IL-8 secretion was strictly controlled as the microbiome metabolites tested here did not induce any IL-8 response, likely to avoid any inadvertent induction of the potent IL-8 dependent immune response in the gut.

All tested metabolites increased the apical secretion of IL-6, whereas 3M-4-TMAB and UDCA increased basolateral secretion as well. It has been suggested that cells stimulated at the apical surface will only secrete IL-6 apically to ensure migrating immune cells are only activated once they have reached the lumen. Therefore, basolateral secretion induced by 3M-4-TMAB and UDCA might be associated with the overactivation of immune cells within the intestinal tissue leading to inflammation and disease (60, 62, 69). In addition, IL-15 can promote survival of IECs during infection and inflammation; however, overexpression on basolateral surfaces can induce apoptosis and T-cell activation leading to disease, depending on the stimulus (70, 71). Our results show that butyrate, 3, 4-TMAB, 4-TMAP, benzoate and LF82 supernatant all increased IL-15 basolateral secretion, whereas L-tryptophan, 3, 4-TMAB, 4-TMAP, UDCA and GCA increased apical secretion. TMA was the only molecule that did not influence IL-15 secretion. Therefore, it can be suggested that the tested molecules, except TMA, have the potential to induce tissue damage via IL-15 signalling. Furthermore, 3M-4-TMAB and 4-TMAP are the only molecules that increased secretion at apical and basolateral surfaces, hence they have the potential to induce extensive inflammation throughout the intestine by accessing the lumen and underlying neighbouring cell (62, 65, 71). As this study measured the quantity of IL-8, IL-6, and IL-15 without observing the effects on immune cells, it cannot be definitively stated whether the secretion of cytokines elicited a pro- or anti-inflammatory effect. However, the intestinal epithelial barrier function was not impaired while increased cytokine secretion was being observed and therefore no damage to the barrier itself was occurring (63, 72).

GCA is a secondary bile acid produced by the colonic microbiome and is reported to be downregulated in IBD patients (73). Previous studies have shown that GCA has an anti-inflammatory effect by inhibiting LPS-induced macrophage recruitment and proinflammatory cytokine secretion, warranting a potential use as an anti-inflammatory treatment (73). This study has shown that exposing macrophages to a lower concentration of GCA (250 nM) significantly increased the number of intracellular LF82 and stimulated the increased secretion of TNF-α after 72 hours of infection. This finding was unexpected as previous work had suggested an anti-inflammatory and antimicrobial effect of GCA in relation to macrophage function (74). However, studies have shown that bile acids can promote expression of the virulence genes, such as flagellin *fliC*, of the *E. coli* LF82 pathogen used in this study (75). Such factors allow its persistence and growth in the gut. Growth curves of *E. coli* LF82 did not show enhanced growth in the presence of GCA, indicating that any fitness advantage is not due to use of GCA as a nutrient source but rather an outcome of the host-pathogen interaction during stress conditions which in turn exacerbates inflammation (76). However, given these findings with GCA and the fact that the exact mechanism resulting in increased intracellular bacterial burden is still to be elucidated, this warrants further investigation if GCA is to be investigated as an anti-inflammatory treatment for CD.

The other molecules tested in this study did not affect bacterial burden within macrophages; however, butyrate, benzoate and 3,4-TMAB, did induce an increased secretion of TNF-α. This was not expected as butyrate has previously been found to attenuate the induced hyperinflammatory responses in macrophages in response to pathogens by decreasing mTOR kinase activity, resulting in downregulation of inflammatory cytokines including TNF-α (44). Furthermore, benzoate has been found to decrease TNF-α release in kidney macrophages by inhibiting NF-κB activity (77).

Taken together these results indicate that specific microbiome-derived small molecules can direct macrophages to be pro-inflammatory by increasing TNF-α secretion or facilitate intracellular bacterial survival. Furthermore, specific molecules can affect IEC function, altering tight junctions, the secretion of cytokines and IEC survival. However most striking is the interplay between these molecules. They were observed to play competing or complementary roles in these assays underlining the delicate balance at play in the intestine and reiterating that disturbance of the microbiome, and the levels of these molecules, can have dramatic effects on intestinal homoeostasis.

## Supplementary Materials

**Figure S1.**
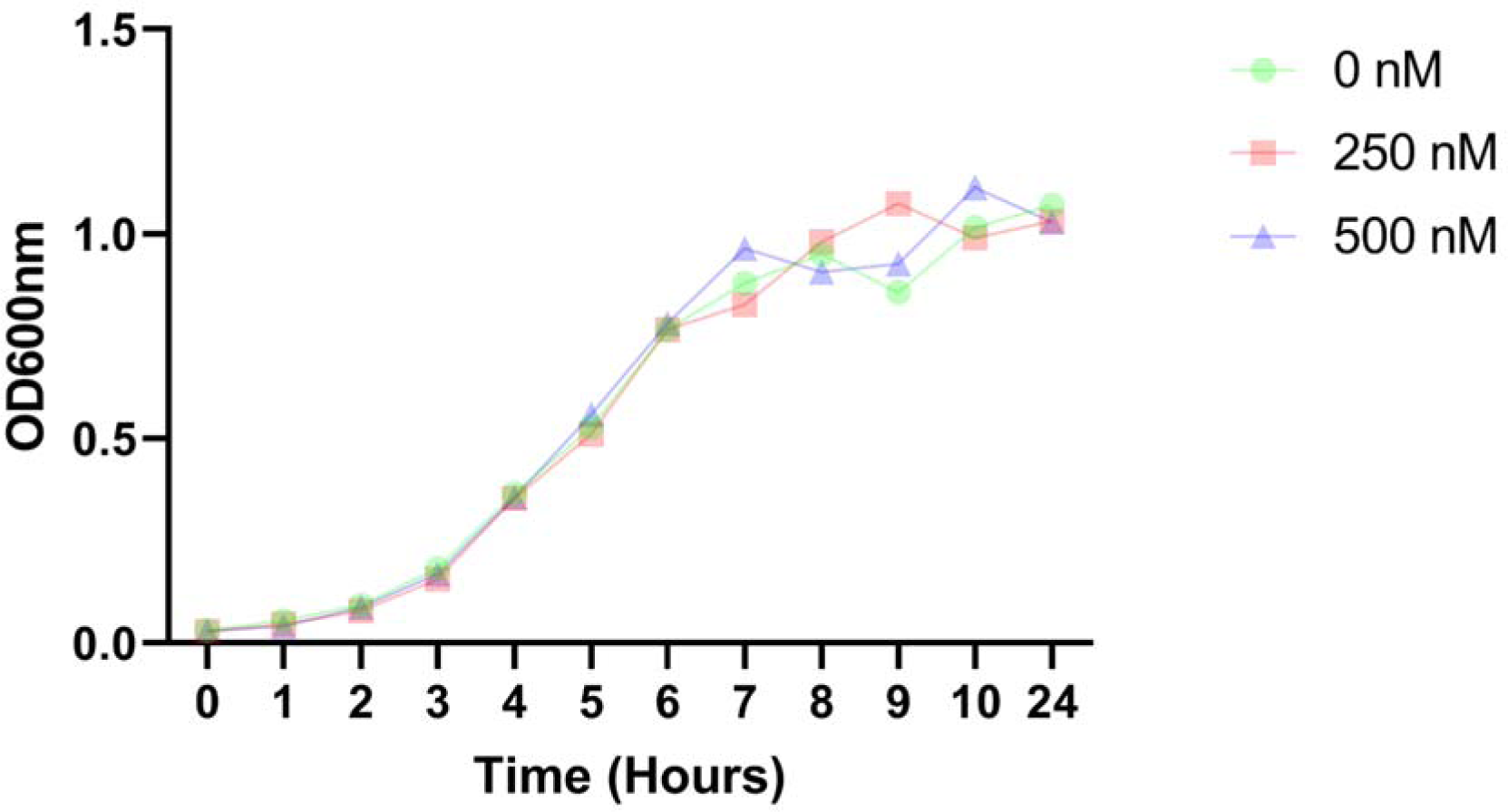
Growth curve of *E. coli* LF82 in GCA supplemented media. Bacteria were cultured in LB and supplemented with 0, 250 or 500 nM GCA and grown at 37°C in a shaken incubator. Results shown as mean of three biological replicates. Two-way ANOVA did not show any significant changes in growth between control condition (0 nM) and the two GCA supplemented conditions.

## Bibliography

1. Clemente JC, Manasson J, Scher JU. The role of the gut microbiome in systemic inflammatory disease. BMJ. 2018;360.

2. Shreiner AB, Kao JY, Young VB. The gut microbiome in health and in disease. Current opinion in gastroenterology. 2015;31(1):69–75.

3. Greenblum S, Turnbaugh PJ, Borenstein E. Metagenomic systems biology of the human gut microbiome reveals topological shifts associated with obesity and inflammatory bowel disease. Proceedings of the National Academy of Sciences of the United States of America. 2012;109(2):594–9.

4. Ko Y, Kariyawasam V, Karnib M, Butcher R, Samuel D, Alrubaie A, et al. Inflammatory Bowel Disease Environmental Risk Factors: A Population-Based Case-Control Study of Middle Eastern Migration to Australia. Clinical Gastroenterology and Hepatology. 2015;13(8):1453–63.e1.

5. Malik TA. Inflammatory Bowel Disease. Historical Perspective, Epidemiology, and Risk Factors. Surgical Clinics of North America. 2015;95(6):1105–22.

6. Caruso R, Lo BC, Núñez G. Host–microbiota interactions in inflammatory bowel disease. Nature Reviews Immunology. 2020;20(7):411–26.

7. Qiu P, Ishimoto T, Fu L, Zhang J, Zhang Z, Liu Y. The Gut Microbiota in Inflammatory Bowel Disease. Frontiers in Cellular and Infection Microbiology. 2022;12:733992-.

8. Salim SAY, Söderholm JD. Importance of disrupted intestinal barrier in inflammatory bowel diseases. Inflammatory Bowel Diseases. 2011;17(1):362–81.

9. Artis D. Epithelial-cell recognition of commensal bacteria and maintenance of immune homeostasis in the gut. Nature Rev Immunol. 2008;8(6):411–20.

10. Bjarnason I, Macpherson A, Hollander D. Intestinal permeability: An overview. Gastroenterology. 1995;108(5):1566–81.

11. Bischoff SC, Barbara G, Buurman W, Ockhuizen T, Schulzke JD, Serino M, et al. Intestinal permeability - a new target for disease prevention and therapy. BMC Gastroenterology. 2014;14(1).

12. Michielan A, D’Incà R. Intestinal Permeability in Inflammatory Bowel Disease: Pathogenesis, Clinical Evaluation, and Therapy of Leaky Gut. Mediators of Inflammation. 2015;2015.

13. Merga Y, Campbell BJ, diseases JMRD, undefined. Mucosal barrier, bacteria and inflammatory bowel disease: possibilities for therapy. kargercomY Merga, BJ Campbell, JM RhodesDigestive diseases, 2014•kargercom.

14. Vindigni SM, Zisman TL, Suskind DL, Damman CJ. The intestinal microbiome, barrier function, and immune system in inflammatory bowel disease: a tripartite pathophysiological circuit with implications for new therapeutic directions. Therapeutic Advances in Gastroenterology. 2016;9(4):606-.

15. Hooper LV, Immunology AJMNR, undefined. Immune adaptations that maintain homeostasis with the intestinal microbiota. naturecomLV Hooper, AJ MacphersonNature Reviews Immunology, 2010•naturecom.

16. Berkes J, Viswanathan VK, Savkovic SD, Hecht G. Intestinal epithelial responses to enteric pathogens: effects on the tight junction barrier, ion transport, and inflammation. Gut. 2003;52(3):439–51.

17. Hooper LV, Wong MH, Thelin A, Hansson L, Falk PG, Gordon JI. Molecular Analysis of Commensal Host-Microbial Relationships in the Intestine. Science. 2001;291(5505):881–4.

18. Zareie M, Riff J, Donato K, McKay DM, Perdue MH, Soderholm JD, et al. Novel effects of the prototype translocating Escherichia coli, strain C25 on intestinal epithelial structure and barrier function. Cellular Microbiology. 2005;7(12):1782–97.

19. Khaloian S, Rath E, Hammoudi N, Gleisinger E, Blutke A, Giesbertz P, et al. Mitochondrial impairment drives intestinal stem cell transition into dysfunctional Paneth cells predicting Crohn’s disease recurrence. Gut. 2020;69(11):1939–51.

20. Goethel A, Croitoru K, Philpott DJ. The interplay between microbes and the immune response in inflammatory bowel disease. The Journal of Physiology. 2018;596(17):3869-.

21. Kawai T, Akira S. Review Toll-like Receptors and Their Crosstalk with Other Innate Receptors in Infection and Immunity. Immunity. 2011;34:637–50.

22. Steinbach EC, Plevy SE. The role of macrophages and dendritic cells in the initiation of inflammation in IBD. Inflammatory bowel diseases. 2014;20(1):166-.

23. Geremia A, Biancheri P, Allan P, Corazza GR, Sabatino AD. Innate and adaptive immunity in inflammatory bowel disease. Autoimmunity Reviews. 2014;13(1):3–10.

24. Hisamatsu T, Kanai T, Mikami Y, Yoneno K, Matsuoka K, Hibi T. Immune aspects of the pathogenesis of inflammatory bowel disease. Pharmacology and Therapeutics. 2013;137(3):283–97.

25. Korta A, Kula J, Gomułka K. The Role of IL-23 in the Pathogenesis and Therapy of Inflammatory Bowel Disease. International Journal of Molecular Sciences. 2023;24(12):10172-.

26. Zhang S, Morgan X, Dogan B, Martin F-P, Strickler S, Oka A, et al. Mucosal metabolites fuel the growth and virulence of E. coli linked to Crohn’s disease. JCI Insight. 2022;7(10).

27. Escribano-Vazquez U, Verstraeten S, Martin R, Chain F, Langella P, Thomas M, Cherbuy C. The commensal Escherichia coli CEC15 reinforces intestinal defences in gnotobiotic mice and is protective in a chronic colitis mouse model. Scientific Reports. 2019;9(1):11431-.

28. Wikoff WR, Anfora AT, Liu J, Schultz PG, Lesley SA, Peters EC, Siuzdak G. Metabolomics analysis reveals large effects of gut microflora on mammalian blood metabolites. Proceedings of the National Academy of Sciences of the United States of America. 2009;106(10):3698–703.

29. Aldars-García L, Gisbert JP, Chaparro M. Metabolomics insights into inflammatory bowel disease: A comprehensive review. Pharmaceuticals. 2021;14(11):1190-.

30. Hulme H, Meikle LM, Strittmatter N, van der Hooft JJJ, Swales J, Bragg RA, et al. Microbiome-derived carnitine mimics as previously unknown mediators of gut-brain axis communication. Science Advances. 2020;6(11):eaax6328–eaax.

31. Pastorelli L, Salvo CD, Mercado J, Vecchi M, Pizarro T. Central Role of the Gut Epithelial Barrier in the Pathogenesis of Chronic Intestinal Inflammation: Lessons Learned from Animal Models and Human Genetics. Frontiers in Immunology2013.

32. Stolfi C, Maresca C, Monteleone G, Laudisi F. Implication of Intestinal Barrier Dysfunction in Gut Dysbiosis and Diseases. Biomedicines2022.

33. Wine E, Ossa JC, Gray-Owen SD, Sherman PM. Adherent-invasive Escherichia coli target the epithelial barrier. Gut Microbes. 2010;1(2):80-.

34. Calzadilla N, Comiskey SM, Dudeja PK, Saksena S, Gill RK, Alrefai WA. Bile acids as inflammatory mediators and modulators of intestinal permeability. Frontiers in Immunology. 2022;13.

35. Peng L, He Z, Chen W, Holzman IR, Lin J. Effects of Butyrate on Intestinal Barrier Function in a Caco-2 Cell Monolayer Model of Intestinal Barrier. Pediatric Research. 2007;61(1):37–41.

36. Scott SA, Fu J, Chang PV. Microbial tryptophan metabolites regulate gut barrier function via the aryl hydrocarbon receptor. Proceedings of the National Academy of Sciences. 2020;117(32):19376–87.

37. Jaworska K, Konop M, Bielinska K, Hutsch T, Dziekiewicz M, Banaszkiewicz A, Ufnal M. Inflammatory bowel disease is associated with increased gut-to-blood penetration of short-chain fatty acids: A new, non-invasive marker of a functional intestinal lesion. Experimental Physiology. 2019;104(8):1226–36.

38. Walczak-Nowicka ŁJ, Herbet M. Sodium Benzoate—Harmfulness and Potential Use in Therapies for Disorders Related to the Nervous System: A Review. Nutrients. 2022;14(7).

39. Campbell HE, Escudier MP, Patel P, Challacombe SJ, Sanderson JD, Lomer MCE. Review article: cinnamon- and benzoate-free diet as a primary treatment for orofacial granulomatosis. Alimentary Pharmacology & Therapeutics. 2011;34(7):687–701.

40. Shi X, Li X, Li X, He Z, Chen X, Song J, et al. Antibacterial Properties of TMA against Escherichia coli and Effect of Temperature and Storage Duration on TMA Content, Lysozyme Activity and Content in Eggs. Foods. 2022;11(4).

41. Chen H, Zhong Q. Antibacterial activity of acidified sodium benzoate against Escherichia coli O157:H7, Salmonella enterica, and Listeria monocytogenes in tryptic soy broth and on cherry tomatoes. International journal of food microbiology. 2018;274:38–44.

42. Lin MH, Khnykin D. Fatty acid transporters in skin development, function and disease. Biochimica et biophysica acta. 2014;1841(3):362-.

43. Deng J, Zhao F, Yu X, Zhao Y, Li D, Shi H, Sun Y. Expression of Aquaporin 4 and Breakdown of the Blood-Brain Barrier after Hypoglycemia-Induced Brain Edema in Rats. PLoS ONE. 2014;9(9).

44. Chen J, Vitetta L. The Role of Butyrate in Attenuating Pathobiont-Induced Hyperinflammation. Immune network. 2020;20(2):e15-e.

45. Zheng L, Kelly CJ, Battista KD, Schaefer R, Lanis JM, Alexeev EE, et al. Microbial-derived Butyrate Promotes Epithelial Barrier Function Through IL-10 Receptor-dependent Repression of Claudin-2. Journal of immunology (Baltimore, Md: 1950). 2017;199(8):2976-.

46. Nunes T, Bernardazzi C, Souza HSD. Cell Death and Inflammatory Bowel Diseases: Apoptosis, Necrosis, and Autophagy in the Intestinal Epithelium. BioMed Research International. 2014;2014.

47. Ruemmele FM, Schwartz S, Seidman EG, Dionne S, Levy E, Lentze MJ. Butyrate induced Caco-2 cell apoptosis is mediated via the mitochondrial pathway. Gut. 2003;52(1):94–100.

48. Pant K, Yadav AK, Gupta P, Islam R, Saraya A, Venugopal SK. Butyrate induces ROS-mediated apoptosis by modulating miR-22/SIRT-1 pathway in hepatic cancer cells. Redox Biology. 2017;12:340–9.

49. Zamaraeva MV, Sabirov RZ, Maeno E, Ando-Akatsuka Y, Bessonova SV, Okada Y. Cells die with increased cytosolic ATP during apoptosis: a bioluminescence study with intracellular luciferase. Cell Death & Differentiation 2005 12:11. 2005;12(11):1390–7.

50. Guerbette T, Boudry G, Lan A. Mitochondrial function in intestinal epithelium homeostasis and modulation in diet-induced obesity. Molecular Metabolism. 2022;63:101546-.

51. Mottawea W, Chiang C-K, Mühlbauer M, Starr AE, Butcher J, Abujamel T, et al. Altered intestinal microbiota–host mitochondria crosstalk in new onset Crohn’s disease. Nature Communications. 2016;7:13419-.

52. Jackson DN, Panopoulos M, Neumann WL, Turner K, Cantarel BL, Thompson-Snipes L, et al. Mitochondrial dysfunction during loss of prohibitin 1 triggers Paneth cell defects and ileitis. Gut. 2020;69(11):1928 LP-38.

53. Nicholls DG. Mitochondrial membrane potential and aging. Aging Cell. 2004;3(1):35–40.

54. Li Y, Yang S, Jin X, Li D, Lu J, Wang X, Wu M. Mitochondria as novel mediators linking gut microbiota to atherosclerosis that is ameliorated by herbal medicine: A review. Frontiers in Pharmacology. 2023;14:52-.

55. Tars K, Leitans J, Kazaks A, Zelencova D, Liepinsh E, Kuka J, et al. Targeting carnitine biosynthesis: discovery of new inhibitors against γ-butyrobetaine hydroxylase. Journal of medicinal chemistry. 2014;57(6):2213–36.

56. Nguyen D, Samson SL, Reddy VT, Gonzalez EV, Sekhar RV. Impaired mitochondrial fatty acid oxidation and insulin resistance in aging: novel protective role of glutathione. Aging cell. 2013;12(3):415–25.

57. Mikó E, Vida A, Kovács T, Ujlaki G, Trencsényi G, Márton J, et al. Lithocholic acid, a bacterial metabolite reduces breast cancer cell proliferation and aggressiveness. Biochimica et Biophysica Acta (BBA) - Bioenergetics. 2018;1859(9):958–74.

58. Kuipers F, Bloks VW, Groen AK. Beyond intestinal soap—bile acids in metabolic control. Nature Reviews Endocrinology 2014 10:8. 2014;10(8):488–98.

59. Xu W, Li T, Gao L, Lenahan C, Zheng J, Yan J, et al. Sodium benzoate attenuates secondary brain injury by inhibiting neuronal apoptosis and reducing mitochondria-mediated oxidative stress in a rat model of intracerebral hemorrhage: Possible involvement of DJ-1/Akt/IKK/NFκB pathway. Frontiers in Molecular Neuroscience. 2019;12:105-.

60. Tapia R, Kralicek SE, Hecht GA. Modulation of epithelial cell polarity by bacterial pathogens. Annals of the New York Academy of Sciences. 2017;1405(1):16-.

61. Gwilt KB, Thiagarajah JR. Membrane Lipids in Epithelial Polarity: Sorting out the PIPs. Frontiers in Cell and Developmental Biology. 2022;10:1124-.

62. Stoops EH, Caplan MJ. Trafficking to the Apical and Basolateral Membranes in Polarized Epithelial Cells. Journal of the American Society of Nephrology: JASN. 2014;25(7):1375-.

63. Andrews C, McLean MH, Durum SK. Cytokine Tuning of Intestinal Epithelial Function. Frontiers in Immunology2018.

64. Huang DB, DuPont HL, Jiang Z-D, Carlin L, Okhuysen PC. Interleukin-8 response in an intestinal HCT-8 cell line infected with enteroaggregative and enterotoxigenic Escherichia coli. Clinical and diagnostic laboratory immunology. 2004;11(3):548–51.

65. Healy LL, Cronin JG, Sheldon IM. Polarized epithelial cells secrete interleukin 6 apically in the bovine endometrium. Biology of Reproduction. 2015;92(6).

66. Mirsepasi-Lauridsen HC, Vallance BA, Krogfelt KA, Petersen AM. Escherichia coli pathobionts associated with inflammatory bowel disease. Clinical Microbiology Reviews. 2019;32(2).

67. Russo TA, Johnson JR. Proposal for a New Inclusive Designation for Extraintestinal Pathogenic Isolates of Escherichia coli: ExPEC. The Journal of Infectious Diseases. 2000;181(5):1753–4.

68. Hernandes RT, Elias WP, Vieira MAM, Gomes TAT. An overview of atypical enteropathogenic Escherichia coli. FEMS Microbiology Letters. 2009;297(2):137–49.

69. Cai Y, Sugimoto C, Arainga M, Alvarez X, Didier ES, Kuroda MJ. In Vivo Characterization of Alveolar and Interstitial Lung Macrophages in Rhesus Macaques: Implications for Understanding Lung Disease in Humans. The Journal of Immunology. 2014;192(6):2821–9.

70. Ebert EC. IL-15 converts human intestinal intraepithelial lymphocytes to CD94+ producers of IFN-γ and IL-10, the latter promoting Fas ligand-mediated cytotoxicity. Immunology. 2005;115(1):118-.

71. Mention JJ, Ahmed MB, Bègue B, Barbe U, Verkarre V, Asnafi V, et al. Interleukin 15: A key to disrupted intraepithelial lymphocyte homeostasis and lymphomagenesis in celiac disease. Gastroenterology. 2003;125(3):730–45.

72. Mahapatro M, Erkert L, Becker C. Cytokine-Mediated Crosstalk between Immune Cells and Epithelial Cells in the Gut. Cells2021.

73. Ge X, Huang S, Ren C, Zhao L. Taurocholic Acid and Glycocholic Acid Inhibit Inflammation and Activate Farnesoid X Receptor Expression in LPS-Stimulated Zebrafish and Macrophages. Molecules. 2023;28(5).

74. Zhang YL, Li ZJ, Gou HZ, Song XJ, Zhang L. The gut microbiota-bile acid axis: A potential therapeutic target for liver fibrosis. Front Cell Infect Microbiol. 2022;12:945368.

75. Pi Y, Wu Y, Zhang X, Lu D, Han D, Zhao J, et al. Gut microbiota-derived ursodeoxycholic acid alleviates low birth weight-induced colonic inflammation by enhancing M2 macrophage polarization. Microbiome 2023 11:1. 2023;11(1):1–21.

76. Delmas J, Gibold L, Faïs T, Batista S, Leremboure M, Sinel C, et al. Metabolic adaptation of adherent-invasive Escherichia coli to exposure to bile salts. Scientific Reports. 2019;9(1).

77. Oshima Y, Wakino S, Kanda T, Tajima T, Itoh T, Uchiyama K, et al. Sodium benzoate attenuates 2,8-dihydroxyadenine nephropathy by inhibiting monocyte/macrophage TNF-α expression. Scientific reports. 2023;13(1):3331-.

